# Tiling mechanisms of the compound eye through geometrical tessellation

**DOI:** 10.1101/2022.01.05.475162

**Authors:** Takashi Hayashi, Takeshi Tomomizu, Takamichi Sushida, Masakazu Akiyama, Shin-Ichiro Ei, Makoto Sato

## Abstract

Tilling patterns are observed in many biological structures. Hexagonal tilling, commonly observed in the compound eyes of wild-type *Drosophila*, is dominant in nature; this dominance can probably be attributed to physical restrictions such as structural robustness, minimal boundary length, and space filling efficiency. Surprisingly, tetragonal tiling patterns are also observed in some *Drosophila* small eye mutants and aquatic crustaceans. Herein, geometrical tessellation is shown to determine the ommatidial tiling patterns. In small eye mutants, the hexagonal pattern is transformed into a tetragonal pattern as the relative positions of neighboring ommatidia are stretched along the dorsal-ventral axis. Hence, the regular distribution of ommatidia and their uniform growth collectively play an essential role in the establishment of tetragonal and hexagonal tiling patterns in compound eyes.

## Introduction

Tiling is a common phenomenon observed in nature. Although a two-dimensional space can be covered by a repetitive array of triangular, tetragonal, pentagonal or hexagonal tiles, evolution seems to favor hexagonal tiling pattern in nature. One plausible reason for this could be that hexagonal tiles are mechanically more stable compared to tetragonal tiles due to short perimeters, dense space filling, and structural rigidity. In biological systems, hexagonal patterns are observed frequently, as exemplified by honeycombs and compound eyes, but this does not necessarily mean that all hexagonal tiles are constructed based on the same principle.

The compound eye is an interesting example of tiling, and is often constructed by alternate vertices of the hexagonal arrays of ommatidia, the optical unit of the compound eye. However, some insects exhibit tetragonal facets (Gillies and S., 1951; Meyer-Rochow, 1971; Stumm-Tegethoff and Dicke, 1974; Xue et al., 2015). Some aquatic crustaceans, such as shrimp and lobsters, have evolved with tetragonal facets (Exner, 1891; Grenacher, 1879; Meyer-Rochow, 1975; Parker, 1891). Mantis shrimp is an insightful example as its compound eye has a tetragonal midband region sandwiched between hexagonal dorsal and ventral hemispheres (Manning et al., 1984; Marshall and Land, 1993). Thus, arthropod compound eyes can have tetragonal as well as hexagonal tiling patterns. This casts doubt on the naive explanation that hexagonal tiles recur in nature because of their mechanical stability. In the present study, we demonstrate that geometrical tessellation determines ommatidial tiling patterns.

## Results

### Tetragonal tiling patterns in small eye mutants

Tiling patterns can be dissected genetically using mutants of the fruit fly *Drosophila melanogaster* (Johnson, 2021). While the wild-type strain develops hexagonal arrays of ommatidia, tetragonal arrays can be produced via diverse genetic manipulations such as the loss-of-function mutant of the *eyeless* (*ey*) gene (Hartman and Hayes, 1971; Ready et al., 1976), ectopic expression of the *fringe* (*fng*) gene (Gibson and Schubiger, 2000), the *outstretched* (*os*) allele of the *unpaired* (*upd*) genes, which lacks the shared eye-specific regulatory element of *upd1* and *upd3* (Tsai and Sun, 2004; Wang et al., 2014), and the *Lobe* mutant (Singh et al., 2011) (Figs. 1A, B, S1). Since these manipulations target different genetic pathways, the mechanism that switches the hexagonal to tetragonal pattern has so far been difficult to elucidate. Here, we demonstrate that the tiling pattern of the compound eye is governed by geometrical as well as genetic and physical mechanisms to establish either hexagonal or tetragonal patterns.

**Fig. 1.**
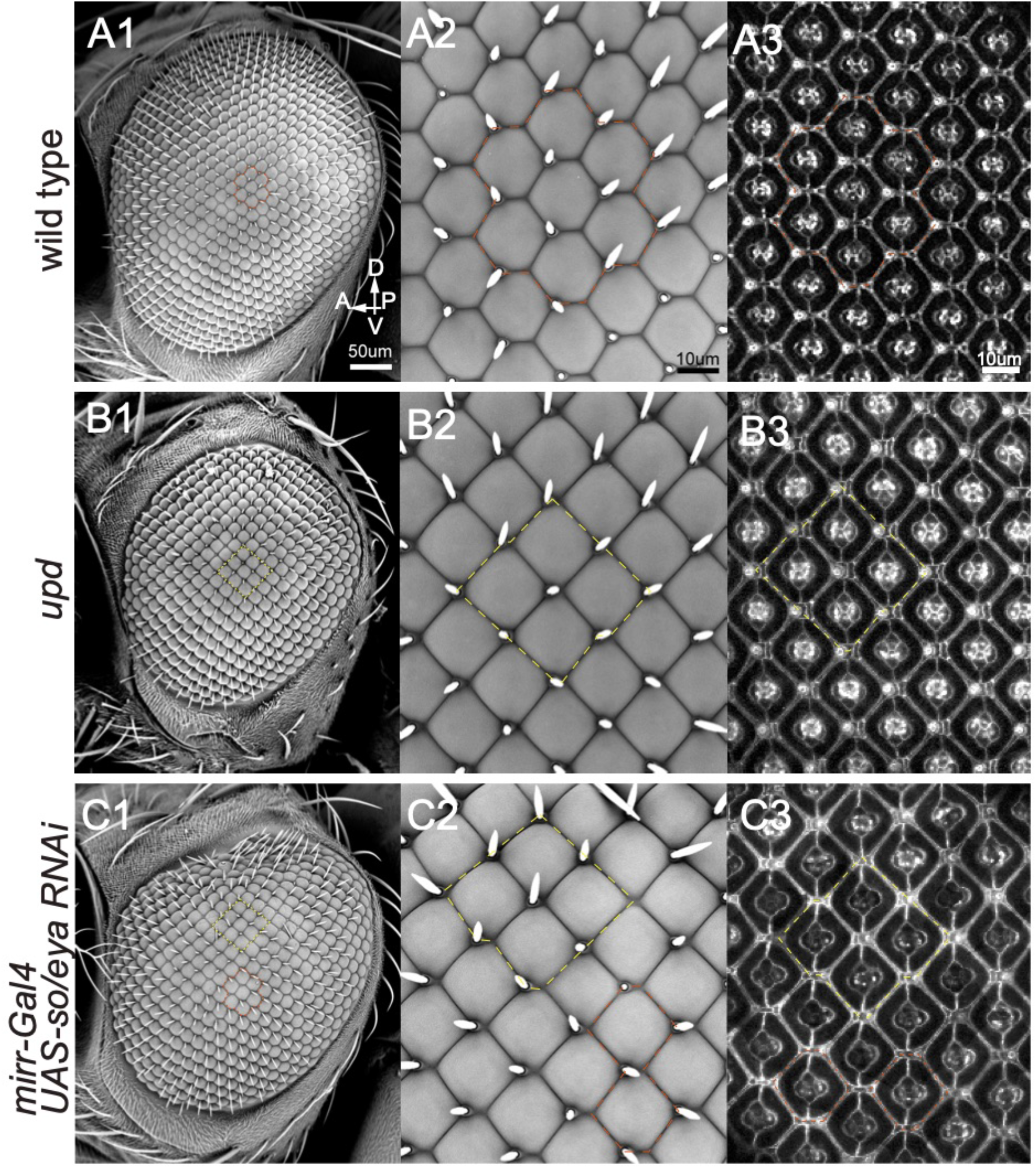
Hexagonal and tetragonal tiling patterns in the fly compound eyes. (A) Hexagonal ommatidial pattern in the wild-type eye (red dotted lines). (B, C) Tetragonal ommatidial patterns (yellow dotted lines) in the *upd^os^* (B) and *mirr-Gal4 UAS-so/eya RNAi* eyes (C). (A1-2, A1-2, A1-2) SEM images. (A3, B3) αCatenin-GFP and (C3) shg-GFP signals at 42 hr APF. Red and yellow dotted lines highlight typical hexagonal and tetragonal ommatidia, respectively.

Interestingly, all the four mutants shared the small-eye phenotype, suggesting that the size of the whole eye determines its tiling pattern. To test whether the small eye size itself was the cause of the tetragonal phenotype, we specifically eliminated the dorsal half of the eye by knocking down the eye specification genes *eyes absent* (*eya*) and *sine oculis* (*so*) under the control of dorsal eye-specific *mirr-Gal4* (Morrison and Halder, 2010; Pignoni et al., 1997). This resulted in small-eye and tetragonal tiling phenotypes (Fig. 1C). In this experiment, the ventral cells were not subjected to any gene manipulation and considered to be genetically wild-type. This experiment confirmed that the size of the whole eye, rather than the autonomous functions of specific genes, is essential for determining whether the tiling pattern is hexagonal or tetragonal.

The tetragonal phenotypes shown above shared the reduced eye size along the dorsal-ventral (DV) axis. In the case of the *upd* mutant, the eye size was reduced along the DV axis because the expression of *upd1* and *upd3*, which activates cell proliferation via Jak/Stat signaling (Tsai and Sun, 2004; Zeidler et al., 1999), was downregulated around the DV border of the eye disc. Overexpression of *fng* may also affect this process (Tsai and Sun, 2004; Zeidler et al., 1999). In contrast, the effect of knocking down *ey* or *Lobe* is mostly limited to the ventral hemisphere (Baker et al., 2018; Chern and Choi, 2002), and is complementary to the elimination of the dorsal hemisphere. Among these mutations, *upd* showed the most uniform ommatidial array with higher regularity and less phenotypic variation (Fig. 1). Therefore, we focused on the *upd* mutant eye as a typical small eye example unless otherwise noted.

### Developmental transition from hexagonal to tetragonal patterns

To identify the origin of the tetragonal facets, the developmental process of the wild-type and the small-eye mutant was evaluated. Five different cell types occur on the surface of the eye: two types of ommatidial cells (cone cells and primary pigment cells) and three types of interommatidial lattice cells (secondary/tertiary pigment cells and bristles; Fig. 2D) (Cagan and Ready, 1989a). The lattice cells specify the ommatidial boundary showing the hexagonal and tetragonal ommatidial shapes at 42 hr after puparium formation (APF; Fig. 2A6, B6) (Cagan and Ready, 1989a). During the early pupal stage at 20 hr APF, the ommatidial tiling pattern was ambiguous and ommatidial spacing was irregular (Fig. 2A1, B1). The tiling pattern became much more apparent and spacing became mostly uniform within a few hours (~24 hr APF, Fig. 2A2, B2).

**Fig. 2.**
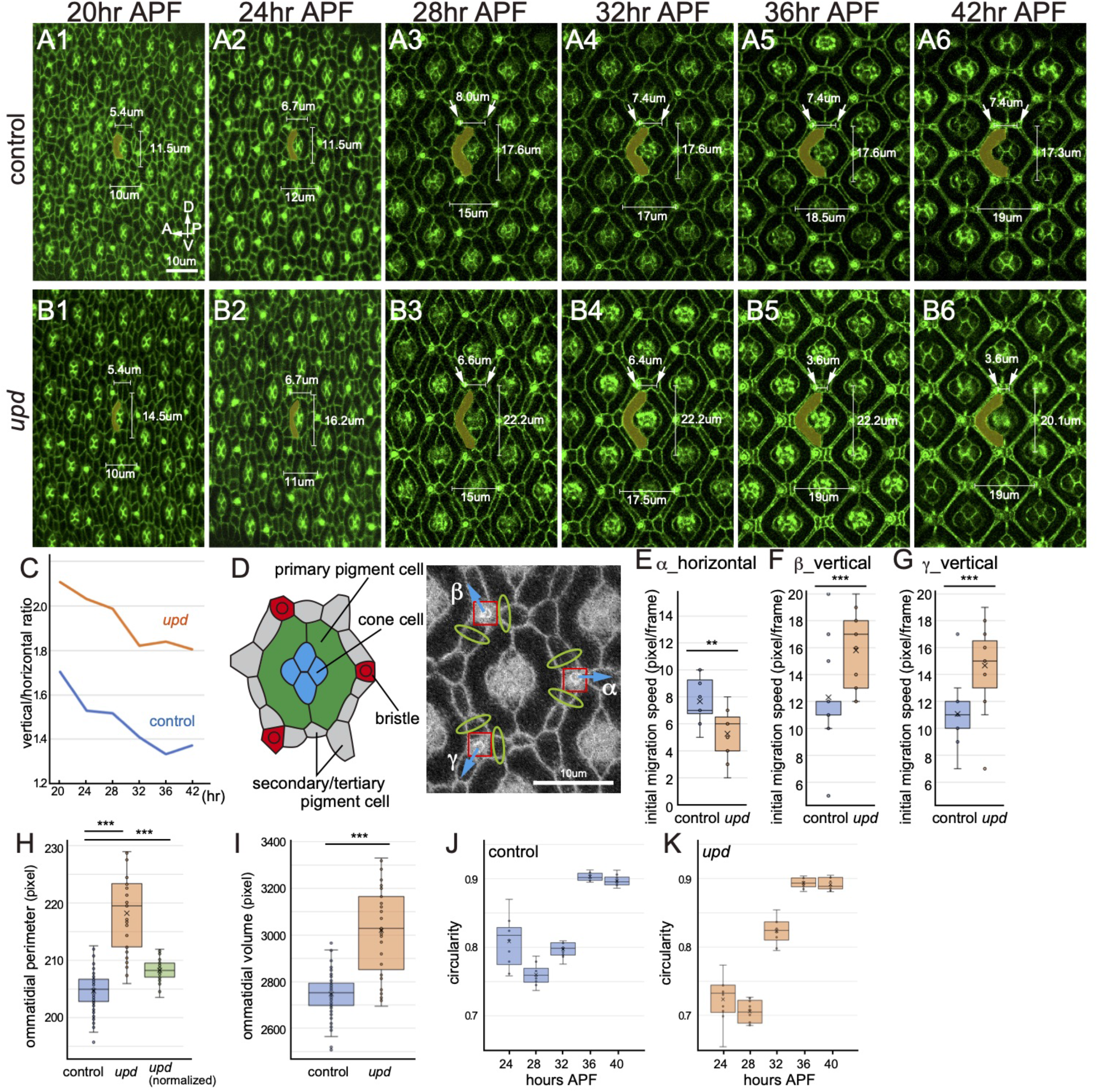
Transition from hexagonal to tetragonal tiling pattern during pupal development. (A, B) Time lapse images from 20 to 42 hr APF (*αCatenin-GFP*). (A) Wild-type eye. (B) *upd* eye. Typical primary pigment cells of the central ommatidia are painted in yellow. Arrows indicate the two corner cells of the dorsal side of the central ommatidia. Lengths of the dorsal side and vertical and horizontal length of the central ommatidia are indicated. (C) The ratio of vertical/horizontal distance between ommatidia during pupal development (n=20). The ommatidial distance was measured based on the centroids of cone cell clusters. (D) Schematics of an ommatidium at 28hr APF. (E-G) Initial migration speeds of a along the horizontal axis (E), and of β (F) and γ (G) along the vertical axis (n=12 and 15 experiments for control and *upd*, respectively). (H, I) Ommatidial perimeter (H) and volume (I) are compared (n=80 and 35 ommatidia for control and *upd*, respectively). Two-sided *t*-test in (E-I; **: p<0.01, ***: p<0.001). (J, K) Ommatidial circularity during pupal development in control (J) and *upd* eyes (K). n=10 ommatidia. Cross, mean; center line, median; box limits, upper and lower quartiles; whiskers, 1.5x interquartile range (E-K).

Interestingly, the tiling process started with the formation of hexagonal patterns in both the wild-type as well as small-eye mutants (28 hr APF, Fig. 2A3, B3). In the *upd* mutant, however, the hexagonal pattern extended vertically along the DV axis. Consistent with this, the ratio of the mean ommatidial distance along the vertical axes (dorsal-ventral) to the horizontal axis (anterior-posterior) was greater in the mutant eye than in the control eye (Fig. 2C). The extended hexagonal phenotype was most likely attributed to the enhanced tension along the DV axis, because the eye reduced in size along the DV axis had to be seamlessly connected with the head capsule, whose size is largely determined by the head size. The elongated hexagon gradually transformed into a tetragon during later stages (28-36 hr APF, Fig. 2B3-5). The vertical/horizontal ratio continuously decreased during this process (Fig. 2C). Meanwhile, both the dorsal and ventral sides of the hexagon continued to shorten, while the width of the middle region continued to increase until the dorsal/ventral sides became very short (arrows in Fig. 2B3-6). Each of the dorsal/ventral sides contained three lattice cells, which were neither eliminated nor they died during this process; instead they ended up getting compressed at the dorsal/ventral vertexes of each tetragon (Fig. 2B5-6).

### Membrane tension is enhanced along the DV axis in the small eye mutant

We speculated that the changes in the ommatidial shape may be caused by membrane tension of the ommatidial cells. Therefore, we measured membrane tension by performing laser ablation experiments at 28hr APF. We focused on three bristle cells, designated as α, β and γ, located at the vertexes of three neighboring ommatidia (red boxes in Fig. 2D). Membrane tension was estimated from the initial migration speeds of the bristle cells immediately after simultaneous ablation of lattice cell membranes adjacent to the bristle cells (green ellipses in Figs. 2D, S2).

We initially focused on the tension along the anterior-posterior (AP) axis because the shortening of the dorsal and ventral sides of ommatidia accompanies the posterior displacement of the bristle cell, α, situated at the posterior vertex of the ommatidium of interest during tetragonization (Fig. 2B3-5, 2D). Note that bristle cell α also constitutes the dorsal and ventral sides of the ommatidia situated posteriorly and may move posteriorly during tetragonization due to the increased tension of the dorsal and ventral lattice cells of the posterior ommatidia. However, its horizontal migration speed following laser ablation was slower in the *upd* ommatidia compared with the control (Figs. 2E, S2). Thus, the membrane tension of the dorsal and ventral lattice cells did not enhance along the AP axis, suggesting that the tetragonization of ommatidia is led by a mechanism that is distinct from the heterogeneous membrane tension observed in the *pten* mutant wing (Bardet et al., 2013).

We next focused on the vertical migration speeds of bristle cells β and γ at 28 hr APF to clarify whether the membrane tension along the DV axis was altered in the *upd* mutant. Consistent with the elongated hexagonal phenotype stretched along the DV axis (Fig. 2B3), their vertical migration speeds were significantly faster in the *upd* mutant (Figs. 2F, G, S2). These results suggested increased tension along the DV axis in the small-eye mutants.

### Tetragonal pattern is not consistent with mechanical optimization

Membrane tension is attributed to actin fibers associated with adherence junctions that accumulates cadherins and catenins, and minimizes the cell perimeter (Del Signore et al., 2018; Honda et al., 1983; Tepass, 1999). The cell composing an ommatidium determines latter’s volume. While the perimeter of tetragonal shapes is not minimized, the cell shape is commonly calculated based on the minimum energy principle that obeys the minimum-perimeter and preferred-volume rule as exemplified by the vertex model (Honda et al., 1983; Honda and Nagai, 2015). We validated these rules on ommatidial patterning. However, both the perimeter and the volume of the *upd* ommatidia were significantly larger than those of the control (Fig. 2H, I). The mean perimeter of the mutant ommatidia normalized by the volume ratio remained larger than that of the control (STAR Methods).

Next, we utilized the vertex model to systematically validate the minimum-perimeter and preferred-volume rule as reported earlier (Honda and Nagai, 2015). Ommatidia show clear structure at the level of adherence junction where the actin fibers produce the membrane tension, and the ommatidial cell shape on the surface of the eye is faithfully reproduced as a two-dimensional structure (Hilgenfeldt et al., 2008; Kafer et al., 2007). Therefore, we utilized the two-dimensional vertex model to determine whether the elongated hexagonal pattern observed in the early *upd* mutant at 28 hr APF could transform into the final tetragonal pattern. However, despite using a wide range of parameters for the perimeter-shortening and volume-conservation terms, the near-square pattern observed in the *upd* ommatidia could not be reproduced (Fig. S3A-C). On the contrary, the near-square pattern of the *upd* mutant at 42hr APF transformed into a hexagonal pattern (Fig. S3D). These results provided a strong evidence that the minimum-perimeter and preferred-volume rule and the vertex model do not support the tetragonal ommatidial pattern.

### The Voronoi diagram recapitulates the hexagonal and tetragonal patterns

The Voronoi diagram, a geometrical partitioning method, is suitable to explain the ommatidial shapes, because it explains the multicellular packing pattern observed in biological system, such as that of the starfish embryo during early development, and was used for a quantitative analysis of the *Drosophila* facet patterning using a two-dimensional model (Fig. 3A; STAR Methods) (Honda et al., 1983; Kim et al., 2016). Therefore, we performed computer simulations based on the Voronoi diagram. Briefly, the center of mass for each cone cell cluster was considered as the mother point. The results of the Voronoi tessellation were compared with the shapes of the ommatidial boundaries extracted from *in vivo* images. Remarkably, the theoretical Voronoi diagram accurately overlapped with the experimental images of the tetragonal mutant eyes as well as those of the hexagonal control eyes at 42 hr APF (Fig. 3C, E, G; STAR Methods), revealing that the ommatidial tiling obeys the Voronoi tessellation.

**Fig. 3.**
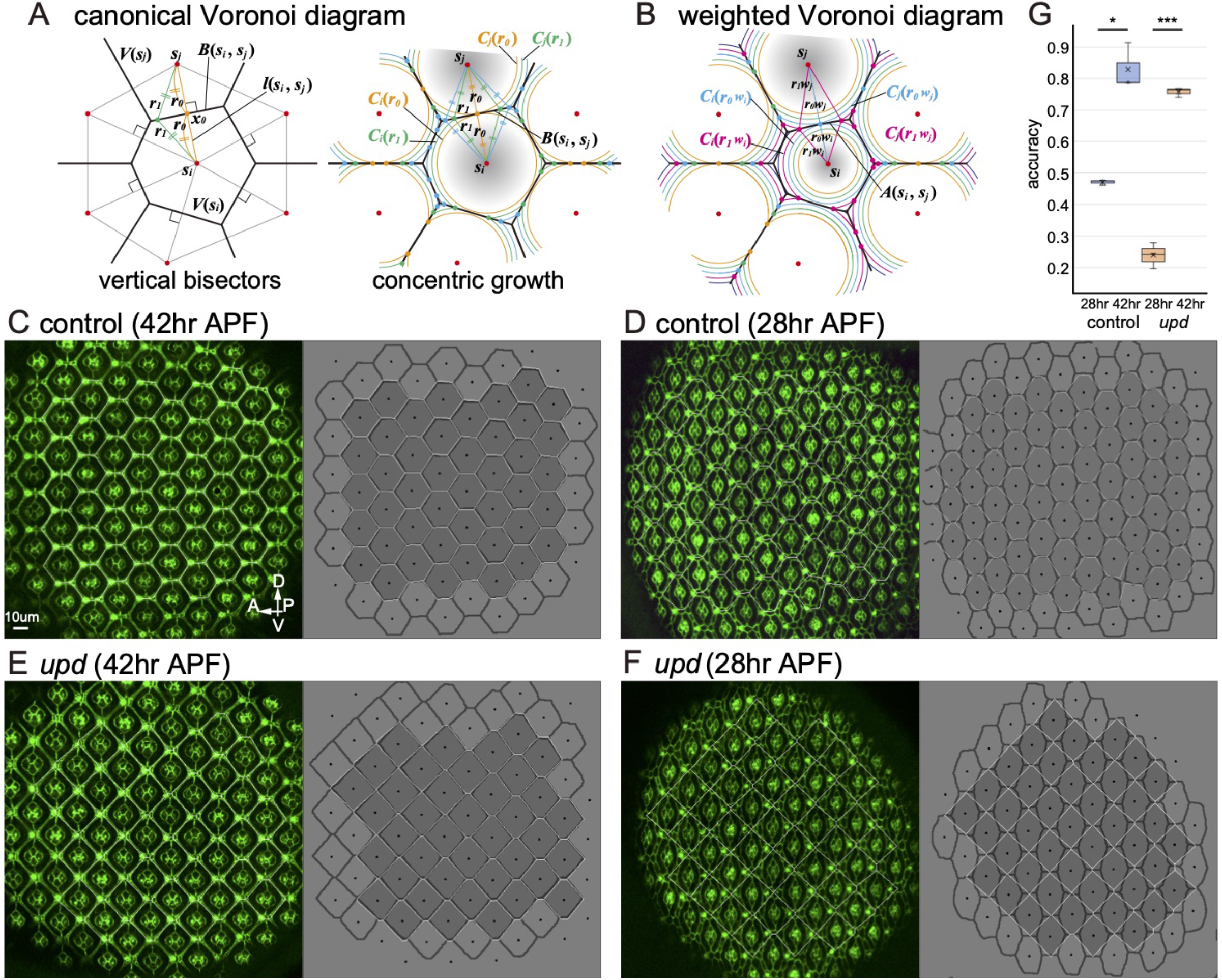
Hexagonal and tetragonal tiling patterns reproduced by the Voronoi diagram. (A) Definition of the canonical Voronoi diagram based on vertical bisectors (left) and concentric growth (right). The concentrically growing circles, *C_i_*(*r*) and *C_j_*(*r*), intersect on the Voronoi edge *B*(***s***_*i*_, ***s***_*j*_), a part of the vertical bisectors of ***s***_*i*_ and ***s***_*j*_. (B) Definition of the weighted Voronoi diagram. The concentrically growing circles with differential growth rates, *C_i_*(*rw_i_*) and *C_j_*(*rw_j_*), intersect on the weighted Voronoi edge *A*(***s***_*i*_, ***s***_*j*_). (C-F) Comparison of the canonical Voronoi diagram (white line) with ommatidial patterns (dark grey line) in control (C, D) and *upd* eyes (E, F). Mature patterns at 42hr (C, E) and immature patterns at 28hr APF (D, F). (G) Quantification of the accuracy of the canonical Voronoi diagram (two-sided *t*-test, *: p<0.05, ***: p<0.001, n=3 eye samples). Cross, mean; center line, median; box limits, upper and lower quartiles; whiskers, 1.5x interquartile range.

Importantly, the Voronoi diagram recapitulated the hexagonal to tetragonal transformation by stretching the DV axis. When the mother points are evenly arranged at the same intervals along the horizontal and vertical axis, the resulting Voronoi pattern becomes hexagonal (Fig. S4). However, when the interval along the vertical axis is gradually increased, the pattern starts to transform into the tetragonal pattern, as the vertical/horizontal ratio becomes close to 2.0. In the small-eye mutant, the vertical/horizontal ratio was between 2.1 and 1.8 during the pupal development, consistent with the near-square patterns observed in the Voronoi diagram (Fig. 2C).

### The Voronoi tessellation is achieved by semi-concentric growth of ommatidia

The next step is to understand how the Voronoi tessellation is achieved *in vivo*. The Voronoi diagram corresponds to the patterns produced by concentric growth of circles; when multiple regions emanating from mother points grow concentrically at the same speed, the resultant pattern recaptures the Voronoi diagram (Figs. 3A, Fig. S5A; STAR Methods). This could explain how ommatidial patterns can be reproduced by the Voronoi diagrams.

Although ommatidia do not have a perfect circular shape, ommatidial cell growth may play a role equivalent to that of the concentric growth. The shape of each ommatidium, outlined by the outer surface of the primary pigment cell pair, was irregular during early stages (Fig. 2A1, B1), but became smooth and circular during later stages (Fig. 2A6, B6). We speculate that the rapid growth of the primary pigment cells may increase the cellular pressure and make the ommatidial shape smoother. This may also result in a semi-concentric growth, centered at the mother point. Indeed, the ommatidial circularity was approximately 0.7 in early stages but increased to approximately 0.9 during later stages (Fig. 2J, K; STAR Methods).

To examine how the growth characteristics of the ommatidia influences the applicability of Voronoi tessellation, we tried to reproduce a more generalized pattern in which the number of ommatidial cells varies among ommatidia. In this scenario, the differential growth rate of each ommatidium should be determined as a function of the number of ommatidial cells it contains, and this should eventually influence the ommatidial tiling pattern.

As a critical test for cell number variability, we chose *Rough eye* (*Roi*), which is a gain-of-function allele of the *amos* gene and exhibits strong variability with fewer and/or extra cone and primary pigment cells (Fig. 4A) (Chanut et al., 2002; Hayashi and Carthew, 2004); our results showed that fewer cells produced smaller ommatidia, while extra cells resulted in larger ommatidia (Figs. 4B, S5D). Notably, the number of primary pigment cells has a much stronger impact on ommatidial size than that of cone cells, suggesting that primary pigment cells are the main determinants of the ommatidial growth rate (Blackie et al., 2020). According to the size of cone and primary pigment cells in the wild-type eyes (Fig. S5C) and the number of these cells in the mutant ommatidium, we estimated the relative growth rate of the *Roi* mutant ommatidia (Fig. 4B; STAR Methods).

**Fig. 4.**
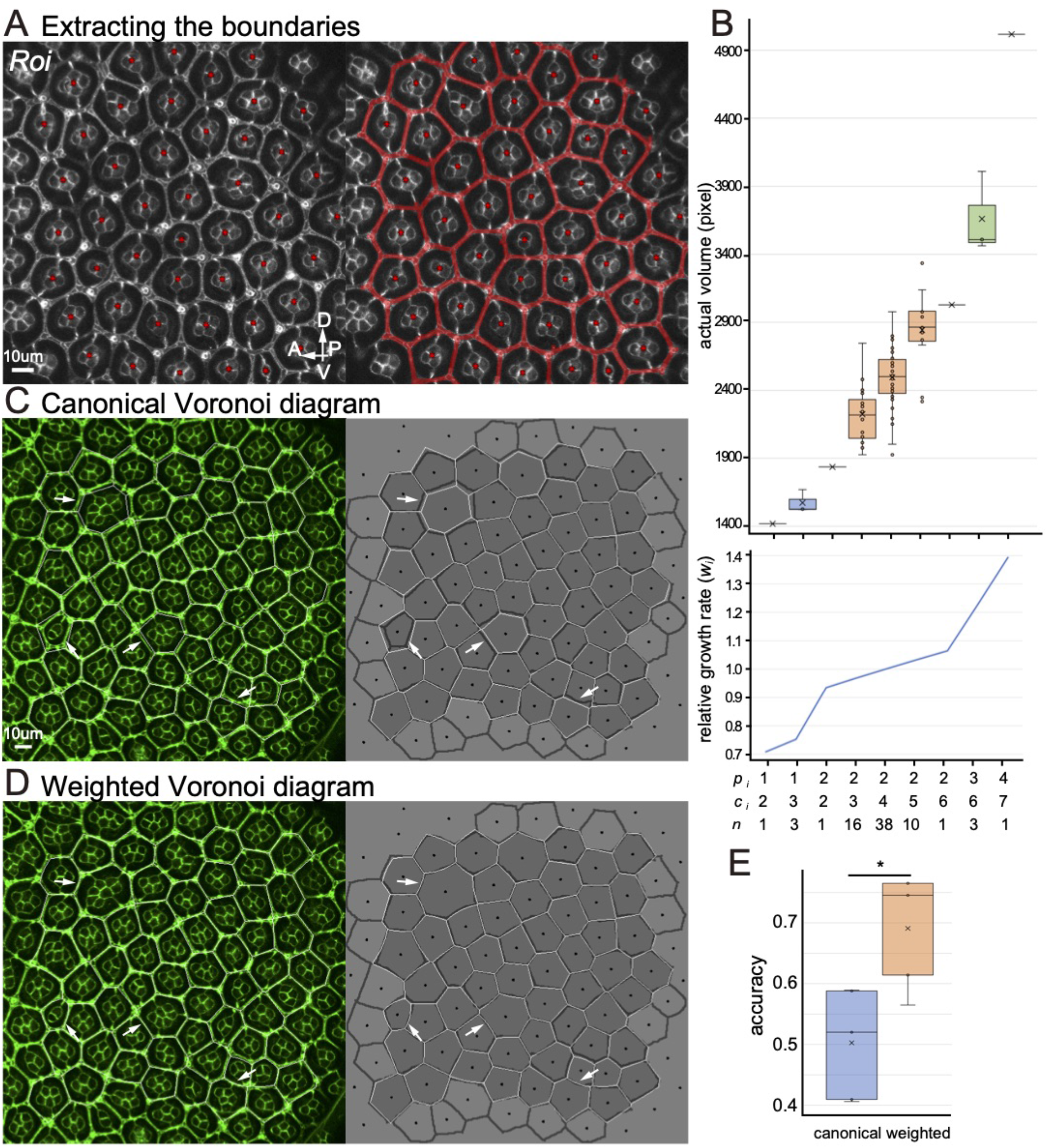
Stochastic tiling patterns reproduced by the Voronoi diagram. (A) *Roi* mutant ommatidia. Red points indicate centers of the cone cell clusters. Red lines indicate the ommatidial boundaries. (B) The ommatidia shown in (A) are classified according to the number of primary pigment cells (*p_i_*) and cone cells (*c_i_*). *n* indicates the number of ommatidia for each class. The actual volume (top) and estimated relative growth rate (bottom, *w_i_*) are plotted. (C, D) Comparison of *Roi* mutant ommatidial patterns (dark grey line) with canonical (white line in C) and weighted Voronoi diagrams (white line in D). Arrows indicate the boundaries between ommatidia with fewer/extra number of the primary pigment cells. (E) Quantification of the accuracy of the canonical and weighted Voronoi diagrams (two-sided *t*-test, *: p<0.05, n=5 eye samples). Cross, mean; center line, median; box limits, upper and lower quartiles; whiskers, 1.5x interquartile range (B, E).

The canonical Voronoi diagram, which is based on the uniform growth rate due to the uniform number of ommatidial cells, did not effectively recapitulate the pattern of extremely small/large ommatidia with fewer/extra primary pigment cells (arrows in Fig. 4C, E). Hence, we utilized the weighted Voronoi diagram, in which the differential growth rate was applied according to the number of ommatidial cells (Fig. 4B, D). This algorithm significantly improved the accuracy, reproducing the irregular tiling patterns at 42 hr APF with high reliability (Fig. 4E). The strong influence of the growth rate on the final pattern suggests that the Voronoi tessellation patterns in compound eyes are achieved by the semi-concentric growth of ommatidial units. Once the arrangement of the mother points and their relative growth rates are known, the Voronoi tessellation can precisely predicts the final ommatidial tiling patterns.

## Discussion

In this study, we showed that the Voronoi diagram could reproduce the final ommatidial patterns. However, it could not reproduce the early patterns. While the control and *upd* ommatidia *in vivo* showed hexagonal patterns at 28 hr APF, the Voronoi diagram predicted hexagonal and tetragonal patterns, respectively (Fig. 3D, F, G). The Voronoi diagram only reproduced the final ommatidial patterns following the semi-concentric ommatidial growth that occurs between 28 and 36 hr APF (Fig. 2J, K) and did not predict the temporal transition of the ommatidial shape during development.

In contrast, using the vertex model, the near-square pattern of the *upd* mutant at 42 hr APF transformed to a hexagonal pattern elongated along the DV axis (Fig. S3D), which is similar to the early ommatidial patterns of the control and *upd* eyes at 28 hr APF (Fig. 3D, F). The changes of ommatidial shape during pupal development were opposite to those observed in the developing starfish embryo, in which the cell shape changed from an early pattern defined by the Voronoi diagram to a late pattern defined by the vertex model, possibly due to the increased membrane tension of the epithelial cells (Honda et al., 1983). Our results may suggest that the shapes of cells and/or ommatidia change to the Voronoi-like and the vertex model-like patterns depending on developmental contexts. An extended vertex model compatible with the semi-concentric ommatidial growth may reproduce the hexagonal to tetragonal transition during ommatidial development.

The hexagonal and tetragonal shapes of ommatidia are believed to have physiological importance. Due to the hexagonal arrangement and the curvature of the *Drosophila* compound eye, photoreceptor neurons R1-6 in an ommatidium receive visual information from different points in space and project to neighboring columns in the brain such that each column receives visual stimuli from the same point in space (Clandinin and Zipursky, 2000; Sato et al., 2013). However, in an aquatic system, the light is not sufficiently focused because of the small difference of refractive indexes between water and lens. The tetragonal arrangement of the shrimp compound eye is essential for its function as a corner reflector (Land, 1976; Vogt, 1975). Thus, the space-filling strategy of ommatidial tiling may be essential for allowing the evolution of these two diverged polygonal structures.

We propose that the final Voronoi pattern depends on the initial distribution pattern of ommatidia. This means that the crystalline arrangement of the compound eye requires a precise repetitive pattern as a pre-requisite, which may be ensured by the periodic appearance of the founder cells (R8) of ommatidia via lateral inhibition during larval development (Cagan and Ready, 1989b; Lubensky et al., 2011; Sato et al., 2013). However, this alone cannot explain the highly precise spacing, since the ommatidial arrangement is not even in the early pupal retina (Fig. 2A1, 2B1). Thus, the semi-concentric ommatidial growth may also play a role in the precise arrangement of ommatidia during pupal development.

It is important to understand the physical laws that govern biological shapes, especially in developmental biology, constructive biology, and regenerative medicine. Our results provide valuable evidence that geometrical patterns based on the semi-concentric ommatidial growth play an important role in biological patterning. As the optical characteristics of biological visual systems have been leveraged in new technologies, such as artificial compound eyes (Glaeser, 2012; Wu et al., 2017), information regarding semi-concentric ommatidial growth may have potential applications in bionics-related research in the future.

## Acknowledgements

We thank R. Carthew, Y. Hiromi, T. Kojima, Y. Tanaka and H. Togashi for critical comments on the manuscript. We thank K. Kimura for supporting the use of SEM. We thank R. Takayama and members of Sato lab for supporting fly work, Bloomington *Drosophila* Stock Center and *Drosophila* Genetic Resource Center, Kyoto for flies. This work was supported by CREST from JST (JPMJCR14D3 to S. E. and M.S.), Grant-in-Aid for Scientific Research (B), (C), Grant-in-Aid for Scientific Research on Innovative Areas and Grant-in-Aid for Early-Career Scientists from MEXT (18K06251 to T.H., 18K13452 to T.S., 15H05857, 19K03611 and 20H05948 to M.A., and 17H03542, 17H05739, 17H05761 and 19H04771 to M.S.), Takeda Science Foundation, Uehara Memorial Foundation and Cooperative Research of ‘Network Joint Research Center for Materials and Devices’ (to M.S.).

## Author Contributions

T.H. and M.S. conceived and designed the experiments. T.H. performed experiments. T.H., T.T. and M.S. acquired, analyzed and interpreted the data. T.S., M.A., S.E. and M.S. formulated the mathematical models. T.H. and M.S. wrote the manuscript.

## Materials and Methods

### Mathematical modeling and numerical simulation

#### Vertex model

##### 0. Intuitive explanation of the vertex model

The adherence junction of ommatidial cells accumulates actin fibers, whose tension would minimize cell perimeter. Since each ommatidial cell has a particular cell size, each ommatidium may have a preferred ommatidial size, too. To calculate the ommatidial shape based on these basic characteristics, we utilized the following vertex dynamics model (hereinafter “vertex model”) (Honda and Nagai, 2015; Honda et al., 2004).

Although an ommatidium of *Drosophila* is a three-dimensional organ composed of multiple cells, the actin fibers at the adherence junction produce the membrane tension that largely determines the ommatidial shape in a two-dimensional plane containing the adherence junction. For the sake of simplicity, we regard a single ommatidium as a hexagon with six vertices in a two-dimensional plane.

Motion of all the vertices were calculated so that the total ommatidial perimeters are minimized and individual ommatidial volumes become closer to the preferred volume. *s* and *k* are the coefficients that determine the influence of ommatidial perimeter and ommatidial volume on the motion of the vertices, respectively.

##### 1. Calculation of the vertex model

In the experimental observations, both the control and *upd* mutant ommatidia show the hexagonal shape at 28 hr APF. In later stage at 42 hr APF, the former changes to the near-regular hexagons and the latter changes to the near-square tetragons (Fig. 2A, B). We asked if the early hexagonal shape becomes tetragons by testing a wide range of parameter settings based on the vertex model. For this purpose, topology data that link the coordinates of the vertices and the identity of each ommatidium was extracted from 512 x 512 pixel binary images in which only ommatidial boundaries are indicated using ImageJ and a custom-made MATLAB code.

In order to obtain the driving equation of the vertices, the following energy function *U* was defined.

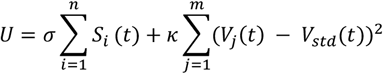

Here, the first term on the right side is the surface energy, which is proportional to the sum of ommatidial perimeters, *S_i_*(*t*). The second term is the area conservation term, which is proportional to a square of the difference of individual ommatidial volume, *V_j_*(*t*), from the preferred volume, *V_std_*(*t*). *S_i_*(*t*) and *V_j_*(*t*) are the length of *i*-th ommatidial perimeter and the volume of, *j*-th ommatidium at time, *t*. *n* and *m* are number of ommatidia. *σ* and *κ* are the positive constants that influence the efficiency of perimeter shortening and area conservation of ommatidia, respectively. Since *U* consists of all vertices 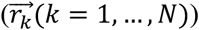, the following vertex driving equations can be obtained by considering the gradient system.

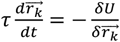

Here, *τ* is a time constant and is set to 1. 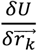 is the functional derivative of *U*.

The calculation was performed using the above gradient system using the Euler method using a custom-made C code. The time step of the calculation was *dt*=1.0 x 10^-4^. The calculation was ended when the rate of change of *U* became sufficiently small 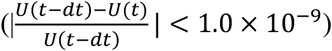. When the volume of an ommatidium became less than half of the initial size, the calculation was stopped because such an abnormal reduction of ommatidia is never observed *in vivo*. The parameters *s* and *k* are in the range of 1.0 x 10^-13^ < *s* < 1.0 x 10^-1^, 1.0 x 10^-13^ < *k* < 1.0 x 10^-8^, respectively. *V_std_*(*t*) is the preferred volume of ommatidia and was defined as follows so that the ommatidial volume changes gradually.

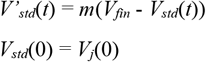

Here, *m* is a positive constant and is set to 1. *V_fin_* is the final volume of ommatidia. From this formula, the preferred volume *V_std_*(*t*) gradually changes from the initial volume at 28 hr APF approaching the final volume *V_fin_*. The mean volume of the control ommatidia at 42 hr APF was 1.20 times larger than that at 28 hr APF (n=3 eye samples). Similarly, the *upd* ommatidia was 1.23 times enlarged at 48 hr APF compared with those at 28 hr APF (n=3 eye samples). Therefore, *V_fin_* = 1.2 x *V_j_* was set in the following calculations.

##### 2. Quantification of the ommatidial shape

In order to quantitatively evaluate the shape of ommatidia, *EdgeRatio* (*ER*) was defined as *ER*=*l_min_*/*l_max_*. Here, *l_max_* and *l_min_* are the the maximum and minimum edge length of an ommatidium, respectively. *ER* is 1 when an ommatidial is composed of equal length edges, such as a regular hexagon. In the case of square ommatidia in the *upd* mutant, it approaches 0, since the dorsal and ventral edges of an ommatidia becomes very short (Fig. 2B6).

The average of *ER* for all the ommatidia is defined as *MeanEdgeRatio* (*MER*), which varies between 0 and 1. The phase diagrams in Fig. S3 indicates *MER* values for a wide range of *s* and *k*. *R_min_* is a parameter set that produces the minimal *MER*. The final shape of the central ommatidium under *R_min_* is shown. When *s* is larger compared with *k*, the ommatidia tend to be abnormally small showing irregular shape, and the calculation was stopped when the volume of an ommatidium became less than half of the initial size.

In the control ommatidia *in vivo, MER* was around 0.7 at 28 and 42 hr APF. In contrast, *MER* was significantly reduced from 0.45 to 0.2 in the *upd* ommatidia (Fig. S3C, n=3 eye samples). In the control simulations using three independent initial conditions at 28 hr APF, the final *MER* value was as large as that of the control at 42 hr APF *in vivo* (Fig. S3A, C). In the *upd* mutant simulations using three initial conditions at 28 hr APF, the final *MER* value was about 0.35, which was significantly larger than that of the *upd* mutant at 42 hr APF *in vivo* (Fig. S3C). Note that ommatidia show the elongated hexagonal shapes even when *MER* is minimized (*MER*=0.30, Fig. S3B).

We also performed the *upd* mutant simulations using the initial condition at 42 hr APF using a parameter set comparable with *R_min_* shown above (*s*=0.05, *k*=1.0×10^-11^). The initial near-square pattern was changed to the elongated hexagonal pattern (Fig. S3D). Note that the dorsal and ventral edges of ommatidia are elongated.

#### Voronoi diagram

##### 0. Intuitive explanation of the Voronoi diagram

When we focus on a pair of mother points ***s***_*i*_ and ***s***_*j*_, the line *l*(***s***_*i*_, ***s***_*j*_) connects the two points (Fig. 3A, left). The line *B*(***s***_*i*_, ***s***_*j*_) is a part of the vertical bisector of *l*(***s***_*i*_, ***s***_*j*_) where the distance from ***s***_*i*_ and ***s***_*j*_ are closer than that from the other mother points, ***s***_*k*_(*k* ≠ *i*,*k* ≠ *j*). Note that any point on *B*(***s***_*i*_, ***s***_*j*_) is equally distant from ***s***_*i*_ and ***s***_*j*_. The concentrically growing circles, *C_i_*(*r*) and *C_j_*(*r*), emanating from ***s***_*i*_ and ***s***_*j*_, respectively, with the same growth rate meet with each other on *B*(***s***_*i*_, ***s***_*j*_), because the radius of the two circles, *r*, indicates the distance from ***s***_*i*_ and ***s***_*j*_ such as *r*_0_ and *r*_1_ (Fig. 3A, right). *r*_0_ is the minimum radius of the two circles, *C_i_*(*r*_0_) and *C_j_*(*r*_0_), that meet with each other at a single point on *B*(***s***_*i*_, ***s***_*j*_) (Fig. 3A, right, orange). *r*_1_ is larger than *r*_0_, and the two circles, *C_i_*(*r*_1_) and *C_j_*(*r*_1_), meet with each other at two points on *B*(***s***_*i*_, ***s***_*j*_) (Fig. 3A, right, green). The set of *B*(***s***_*i*_, ***s***_*j*_) between all of the mother points constitutes the canonical Voronoi diagram, and *B*(***s***_*i*_, ***s***_*j*_) is called the Voronoi edge.

When the concentrically growing circles, *C_i_*(*rw_i_*) and *C_j_*(*rw_j_*), emanating from ***s***_*i*_ and ***s***_*j*_ with the differential growth rates, *w_i_* and *w_j_*, respectively, meet with each other, the intersection points of the two circles constitute the curved boundary *A*(***s***_*i*_, ***s***_*j*_) (Fig. 3B), where the weighted distance from ***s***_*i*_ and ***s***_*j*_, *rw_i_* and *rw_j_*, respectively, are closer than that from the other mother points, ***s***_*k*_(*k* ≠ *i,k* ≠ *j*). The two circles, *C_i_*(*r*_0_*w_i_*) and *C_j_*(*r*_0_*wj*), meet with each other at a single point on *A*(***s***_*i*_, ***s***_*j*_) (Fig. 3B, blue). *r*_1_ is larger than *r*_0_, and the two circles, *C_i_*(*r*_1_*w_i_*) and *C_j_*(*r*_1_*w_j_*), meet with each other at two points on *A*(***s***_*i*_, ***s***_*j*_) (Fig. 3B, magenta). The set of *A*(***s***_*i*_, ***s***_*j*_) between all of the mother points constitutes the weighted Voronoi diagram, and *A*(***s***_*i*_, ***s***_*j*_) is called the weighted Voronoi edge.

##### 1. Definition of the canonical Voronoi diagram

We recall a mathematical definition of the canonical Voronoi diagram with a finite point set on the two-dimensional plane 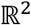. Let 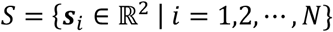 be a point set with *N* elements. Each element ***s***_*i*_ ∈ *S* is called the mother point (Fig. S5A). A canonical Voronoi region with the mother point ***s***_*i*_ is given as follows.

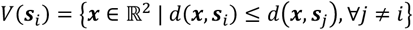

where *d*(***x***, ***y***) denotes the Euclidean distance of 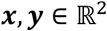. Moreover, the family of Voronoi regions *V*(***s***_*i*_) is given by 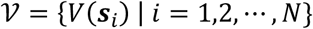, which is called the canonical Voronoi diagram with the point set *S*. Note that the boundary of a Voronoi region is constructed by vertical bisectors of the line segments *l*(***s***_*i*_, ***s***_*j*_) (Fig. 3A, left). If two Voronoi regions *V*(***s***_*i*_) and *V*(***s***_*j*_) are adjacent, the intersection *V*(***s***_*i*_) ⋂ *V*(***s***_*j*_) is a part of the bisector of *l*(***s***_*i*_, ***s***_*j*_), and *V*(***s***_*i*_) ⋂ *V*(***s***_*j*_) is the Voronoi edge *B*(***s***_*i*_, ***s***_*j*_).

Next, we consider the relationship between the Voronoi diagram and the concentric growth of circles at the two mother points ***s***_*i*_ and ***s***_*j*_. For *r* >0, let 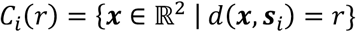 be a circle of the radius *r* with the mother point ***s***_*i*_ ∈ *S* (Fig. S5A). Here we assume that *V*(***s***_*i*_) and *V*(***s***_*j*_) are adjacent. Then, we have *V*(***s***_*i*_) ⋂ *V*(***s***_*j*_) ⊂ *B*(***s***_*i*_, ***s***_*j*_) and there exists *r* = *r*_0_ > 0 such that two circles *C_i_*(*r*_0_) and *C_j_*(*r*_0_) contact with a single point ***x*_0_**. Here, *x*_0_ = *C_i_*(*r*_0_) ⋂ *C_j_*(*r*_0_) and *d*(***x***_0_, ***s***_*i*_) = *d*(***x***_0_, ***s***_*j*_). Therefore, we have ***x***_0_ ∈ *V*(***s***_*i*_) ⋂ *V*(***s***_*j*_) because the distance between ***x***_0_ and ***s***_*i*_, ***s***_*j*_ is the minimum for ***x*** that satisfies the condition *d*(***x***, ***s***_*i*_) = *d*(***x***, ***s***_*j*_). Moreover, the intersection *C_i_*(*r*_1_) ⋂ *C_j_*(*r*_1_) for *r*_1_ > *r*_0_ has two points which lie on the bisector of *l*(***s***_*i*_, ***s***_*j*_) (Fig. S5A). Therefore, the set {*C_i_*(*r*) ⋂ *C_j_*(*r*) | *r* ≥ *r*_0_} is equal to the bisector of *l*(***s***_*i*_, ***s***_*j*_), and hence from the definition of canonical Voronoi region, a set of intersections obtained by the concentric growth of two circles emanating from the two mother points, ***s***_*i*_ and ***s***_*j*_ is equivalent to a boundary of the two Voronoi regions, *V*(***s***_*i*_) and *V*(***s***_*j*_).

##### 2. Definition of the weighted Voronoi diagram

Here, we introduce the weighted Voronoi diagram with the point set *S*. Let *w_i_* >0 be the weight for each mother point ***s***_*i*_ ∈ *S* (Fig. S5B). A weighted Voronoi region with the mother point ***s***_*i*_ is given as follows.

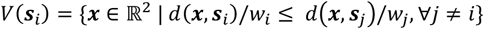

Note that in the above definition, the distance function is modified from the canonical Voronoi region. Therefore, the boundary of weighted Voronoi regions is constructed by an arc (Fig. S5B). If *V*(***s***_*i*_) and *V*(***s***_*j*_) are adjacent, *V*(***s***_*i*_) ⋂ *V*(***s***_*j*_) is a part of the circle defined by 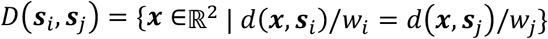 and *V*(***s***_*i*_) ⋂ *V*(***s***_*j*_) is the Voronoi edge *A*(***s***_*i*_, ***s***_*j*_).

Similar to the canonical Voronoi diagram, a set of intersections obtained by the concentric growth of the two circles with differential growth rates, *w_i_* and *w_j_*, is equivalent to a boundary of two weighted Voronoi regions, *V*(***s***_*i*_) and *V*(***s***_*j*_). Here, we consider the modified circle 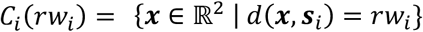 with the radius *rw_i_* of the mother point *s_i_* ∈ *S* for *r* >0 (Fig. S5B). Here, we assume that the two weighted Voronoi regions *V*(***s***_*i*_) and *V*(***s***_*j*_) are adjacent. Then, the set *V*(***s***_*i*_) ⋂ *V*(***s***_*j*_) is contained in the circle *D*(***s***_*i*_, ***s***_*j*_). Also, there exists *r* = *r*_0_ > 0 such that the intersection *C_i_*(*r_0_*w_i_*) ⋂ *C*_j_*(*r*_0_*w_j_*) consists of a single point ***x***_0_ (Fig. S5B). Here, we have ***x***_0_ ∈ *V*(***s***_*i*_) ⋂ *V*(***s***_*j*_) because *d*(***x***, ***s***_*i*_)/*w_i_* = *d*(***x***, ***s***_*j*_)/*w_j_* holds. Moreover, the intersection *C_i_*(*r*_1_*w_i_*) ⋂ *C_j_*(*r*_1_*w_j_*) for *r*_1_ > *r*_0_ has two points which lie on the circle *A*(***s***_*i*_, ***s***_*j*_). Hence, the set {*C_i_*(*rw_i_*) ⋂ *C_j_*(*rw_j_*) | *r* ≥ *r*_0_} is equal to the circle *D*(***s***_*i*_, ***s***_*j*_).

##### 3. Quantification of the overlap between Voronoi diagram and Experimental data

Computer simulations based on the canonical and weighted Voronoi diagrams were performed using Mathematica. The results were compared with the experimental data using the following 512 x 512 pixel binary images prepared using ImageJ.

a. Original gray scale images of the developing ommatidia.
b. Binary image in which only cone cells are painted.
c. Binary images in which only ommatidial boundaries are indicated by three-pixel wide lines.

First, the weight of the concentric growth of each ommatidium was calculated according to the number of cone and primary pigment cells (a). Let *c_i_* and *p_i_* be the number of cone and primary pigment cells in the *i*-th ommatidium, respectively, whose growth rate *R_i_* is proportional to 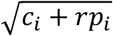, where the value *r*=5.76 indicates the average size of the primary pigment cell area relative to the cone cell area in the wild-type *Drosophila* compound eyes (Fig. S5C). Let 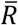 be an average of *R_i_* and define the weight of the growth of the *i*-th ommatidium by 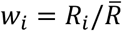 (Fig. 4B). In the case of the canonical Voronoi diagram, *w_i_* =1.

Second, the coordinates of the centroids of cone cell clusters were obtained from (b) using ImageJ (Center of Mass in Measure function). A mother point ***s***_*i*_ is given as the centroid of a cone cell cluster in the *i*-th ommatidium.

The canonical or weighted Voronoi diagram was drawn according to the weight *w_i_* and the mother point ***s***_*i*_, and the result was overlaid on a 512 x 512 pixel binary image as a set of one-pixel wide lines (d), whose overlap with the experimental data was quantified comparing with (c). The Voronoi edges are represented by the pixel value 1 in (c) and (d). Focusing on the position in (d) with the pixel value 1, the pixels whose values are also 1 in (c) is defined as the overlapping pixels. Accuracy of the Voronoi diagram is defined as follows:

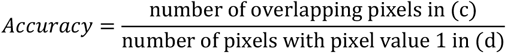

##### 4. Voronoi patterns with different initial conditions

Voronoi diagrams of evenly arranged mother points were drawn using ‘voronoi’ function of MATLAB (Fig. S4).

### Fly strains

Fly strains were maintained on standard *Drosophila* medium at 25°C. *white^1118^* was used as the wild type strain. Mutant strains used were: *upd^os1^* (BDSC#996), *ey^R^* (BDSC#4229), *Lobe^4^* (BDSC#320), *Roi* (BDSC#4105), *mirr-Gal4* (BDSC#29650), *c311-Gal4*, (BDSC#5937), *UAS-fng* (BDSC#5831), *a-Catenin-GFP* (BDSC#59405) and *shg-GFP* (BDSC#58471). The *UAS-so/eya* RNAi strain was generated as described below.

### Generation of *UAS-so/eya* RNAi strain

The UAS-*so/eya* RNAi vector containing both the *eya* and *so* shRNAi sequences was constructed essentially as described (Ni et al., 2011). The *so* shRNAi sequence was cloned between the BsiWI and EcoRI sites and the *eya* shRNA sequence was cloned between the NheI and NotI sites of the pWALIUM20 plasmid. The oligonucleotide sequences used are as follows (the 21bp RNAi sequences are capitalized):

so-Fwd:
gtacgagtTGGTTCAAGAACCGACGACAAtagttatattcaagcataTTGTCGTCGGTTCTTGAACCAgcg
so-Rev:
aattcgcTGGTTCAAGAACCGACGACAAtatgcttgaatataactaTTGTCGTCGGTTCTTGAACCAactc
eya-Fwd:
ctagcagtCAGCAGTTTCTCCACATATCAtagttatattcaagcataTGATATGTGGAGAAACTGCTGgcgc
eya-Rev:
ggccgcgcCAGCAGTTTCTCCACATATCAtatgcttgaatataactaTGATATGTGGAGAAACTGCTGactg

The vector was introduced into the fly genome at the attP40 landing site on the second chromosome.

### Live imaging and laser ablation

For live imaging of the pupal retina, *shg-GFP* (Fig. 1C3) or *a-Catenin-GFP* (all others) was used as the source of GFP. A pupa was sticked on a glass slide with UV curable adhesive and the eye was exposed by removing the pupal case of the head part. The pupa was then surrounded with silicone grease, and a small amount of water was put on the eye and the pupa was mounted with a cover slip. The specimen was analyzed with LSM880NLO (Zeiss) confocal system equipped with Chameleon Vision II pulse laser (COHERENT).

For laser ablation experiment, the retina being live-imaged was irradiated with 700nm two photon laser (30% output) restricted to the six areas that intersect the membrane of the lattice cells as indicated in Fig. 2D (green ellipses). Confocal images were collected every four seconds prior to and following irradiation. One pixel is 0.104mm in our setting.

The migration of bristle cells that belong to the ommatidium of interest were quantified using a custom-made MATLAB code. Three bristle cells designated as α, β, γ (Fig. 2D) were manually selected by 30 x 30 pixel boxes using the image prior to the laser ablation (0 second, Fig. S2A, C). In the following frames, 30 x 30 pixel areas that are most similar to the bristle images of the previous frame were searched among 100 x 100 pixel areas nearby the previous bristle locations. This process was repeated to obtain the time course of the coordinates of the three bristle cells following laser ablation. The time differences of the bristle locations along the horizontal and vertical axes were plotted as migration speed (Fig. 2B, D). Since α, β and γ always moved posteriorly, anterior-dorsally and anterior-ventrally following laser ablation, respectively (blue arrows in Figs. 2D, S2A, C), absolute values of the migration speeds along the horizontal and vertical axes were considered. The migration speeds immediately after laser ablation (4 second) are the initial migration speeds (Figs. 2E-G, S2E-G).

### Quantification of ommatidial shape

Binary images of ommatidial boundaries were prepared from the original gray scale images using ImageJ. Volume (*S*) and perimeter (*L*) of each ommatidium were quantified using Measure function of ImageJ. Perimeter of the *upd* ommatidia (*L_upd_*), average volumes of the control and *upd* ommatidia (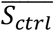 and 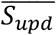, respectively). Normalized perimeter of the *upd* ommatidia was calculated as 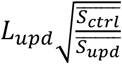 (Fig. 2H). Circularity of ommatidia was calculated as 4π*S* /*L*^2^ (Fig. 2J, K).

### Scanning electron microscopy

Tiling patterns of the compound eyes were examined using a scanning electron microscope (SEM) as described (Kimura et al., 2020). Eyes were fixed with 3.7% formaldehyde and serially dehydrated with ethanol. After dehydration, ethanol was replaced with tert-butyl alcohol. The samples were then frozen at 4 °C and further dehydrated by freeze–drying (VFD-21S, Vacuum Device Inc). Specimens were then subjected to gold coating by an ion coater (Neo coater MP-19010 NCTR, JEOL), and observed with SEM (TM3000 Miniscope, Hitachi).

### Statistics and Reproducibility

For quantification and statistical analysis, distinct eye samples were measured and analyzed. Welch’s t-test was used for the statistical tests. Image intensities were not artificially processed except as otherwise noted. When statistics were not applicable, experiments were independently repeated at least three times with similar results.

