## Supplementary Figures for "Tiling mechanisms of the compound eye through geometrical tessellation"

Fig. S1

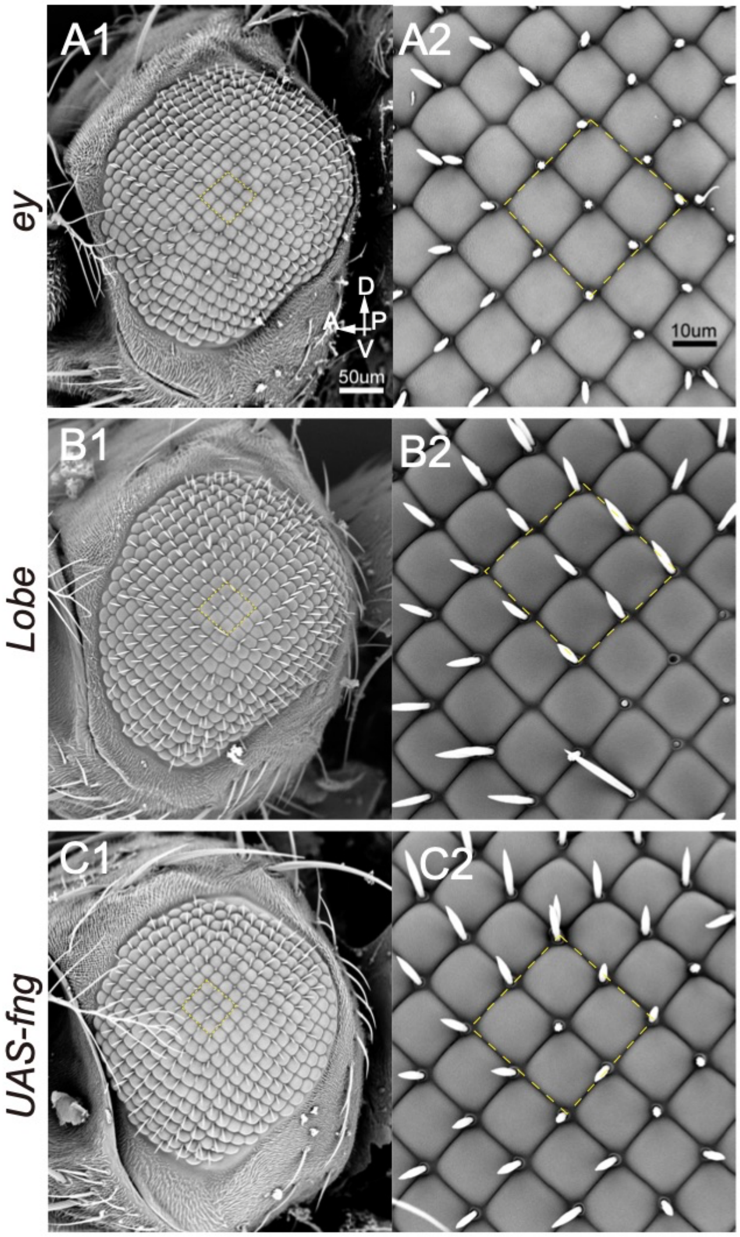


**Fig. S1. Tetragonal tiling patterns observed in mutant eyes.**

SEM images of (A) *ey$^{R}$*, (B) *Lobe$^{4}$* , and (C) *fng* overexpression (*c311-Gal4 UAS-fng*) flies. (A1, B1, C1) Low and (A2, B2, C2) high magnification pictures. Tetragonal patterns are highlighted (yellow dotted lines).

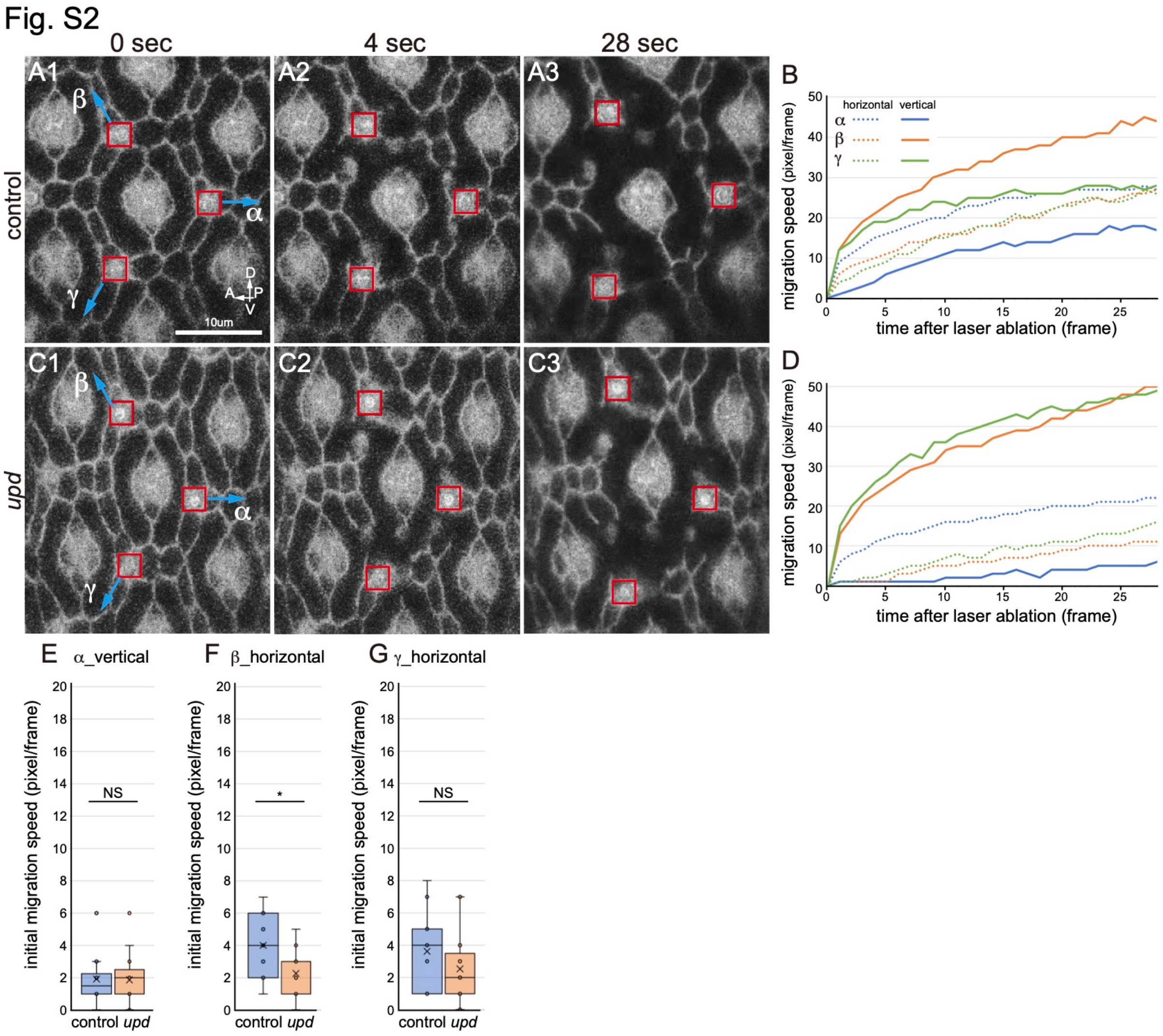


**Fig. S2. Initial migration speed of bristle cells following laser ablation.**

(A, C) Live images of ommatidia at 28 hr APF immediately before laser ablation (A1, C1), and four (A2, C2) and 28 seconds (A3, C3) after laser ablation. The movement of three bristle cells, α, β, γ (red boxes) toward the directions indicated by blue arrows were tracked. (B, D) Time courses of migration speeds of α (blue), β (orange), γ (green) along the vertical and horizontal axes are plotted by regular and dotted lines, respectively. The results of control (A, B) and *upd* ommatidia (C, D) are shown. (E-G) The initial migration speeds of α along the vertical axis (E), and of β (F) and γ (G) along the horizontal axes are compared in the control and *upd* ommatidia (two-sided *t*-test, *: $p<0.05$, **: $p<0.01$, ***: $p<0.001$, n=12 and 15 experiments). Cross, mean; center line, median; box limits, upper and lower quartiles; whiskers, 1.5x interquartile range.

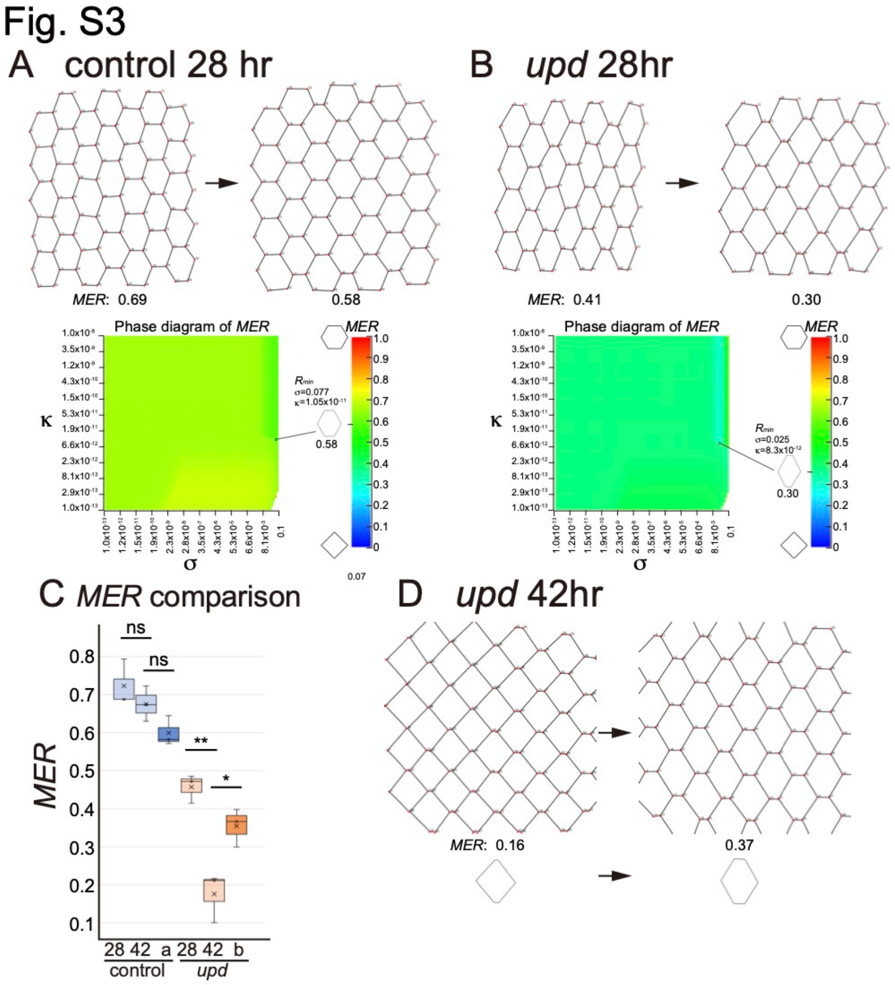


**Fig. S3. Vertex model does not reproduce the square patterns.**

(A, B) Results of computer simulations based on the vertex model. Changes of the ommatidial shapes from the initial to the final states are shown in the upper part. The coordinates of the vertices extracted from the control ommatidia at 28 hr APF (A) and *upd* ommatidia at 28 hr (B) were used as the initial conditions. The phase diagrams in the lower part plot *MER* values in the range of $1.0 \times 10^{-13} < s < 0.1$ and $1.0 \times 10^{-13} < k < 1.0 \times 10^{-8}$. $s$ and $k$ are numerically controlled by the geometric sequences of the common ratios, 1.3219 and 1.1233, respectively (n=100). $R_{min}$ indicates the parameter set that produces the minimal *MER* value. The final shape of the central ommatidium under $R_{min}$ is shown. (C) *MER* values extracted from *in vivo* images of the control and *upd* ommatidia at 28 and 48 hr APF and those of the results of the vertex model simulations in (A) and (B) are compared (two-sided *t*-test, *: $p<0.05$, **: $p<0.01$, n=3 samples). Cross, mean; center line, median; box limits, upper and lower quartiles; whiskers, 1.5x interquartile range. (D) Result of the vertex model simulation using the initial condition of *upd* ommatidia at 42 hr (top). Shape of the central ommatidia (bottom). $s = 0.05$ and $k = 1.0 \times 10^{-11}$.

Fig. S4

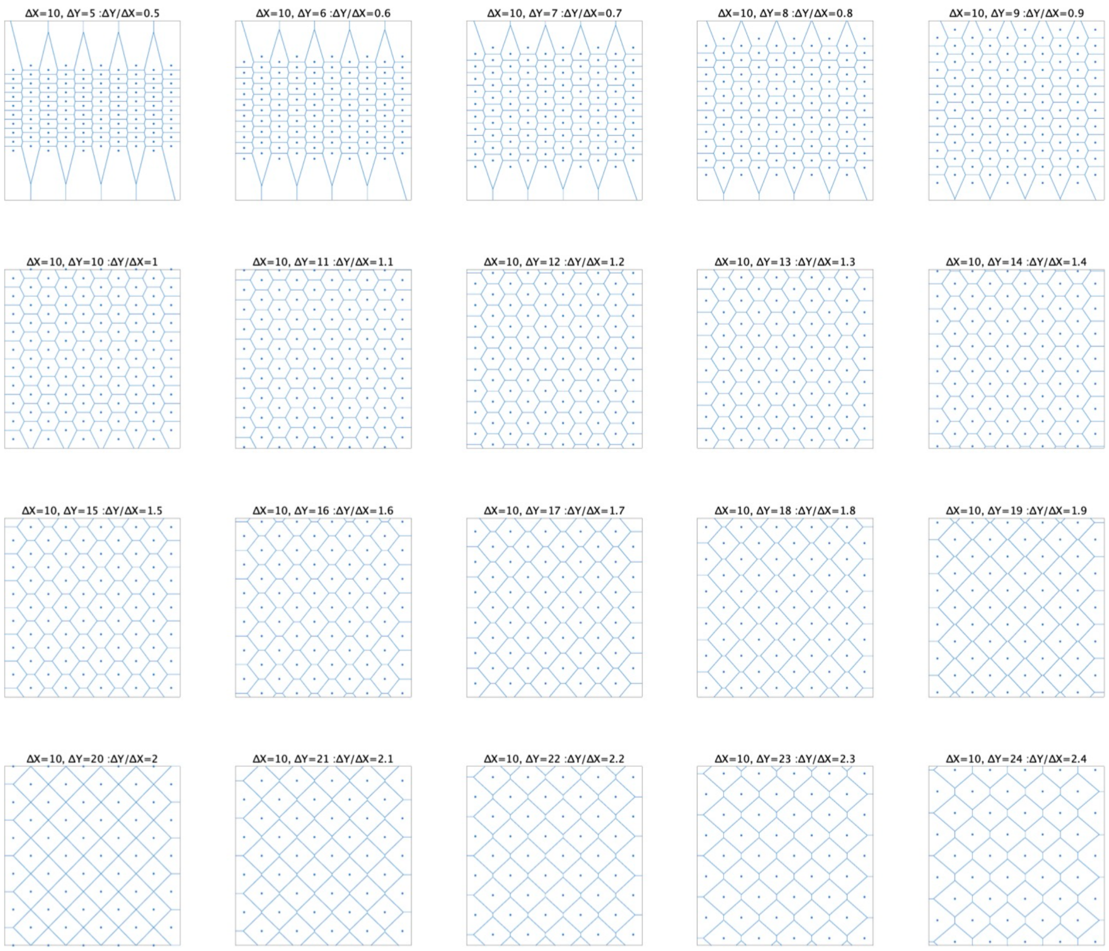


**Fig. S4. Voronoi patterns with different initial conditions.**

Voronoi diagrams of evenly arranged mother points. Horizontal interval of the mother points ($\Delta X$) is always 10, while vertical interval $\Delta Y$) is changed from 5 to 24 so that the vertical/horizontal ratio ($\Delta Y/\Delta X$) varies between 0.5 to 2.4.

Fig. S5

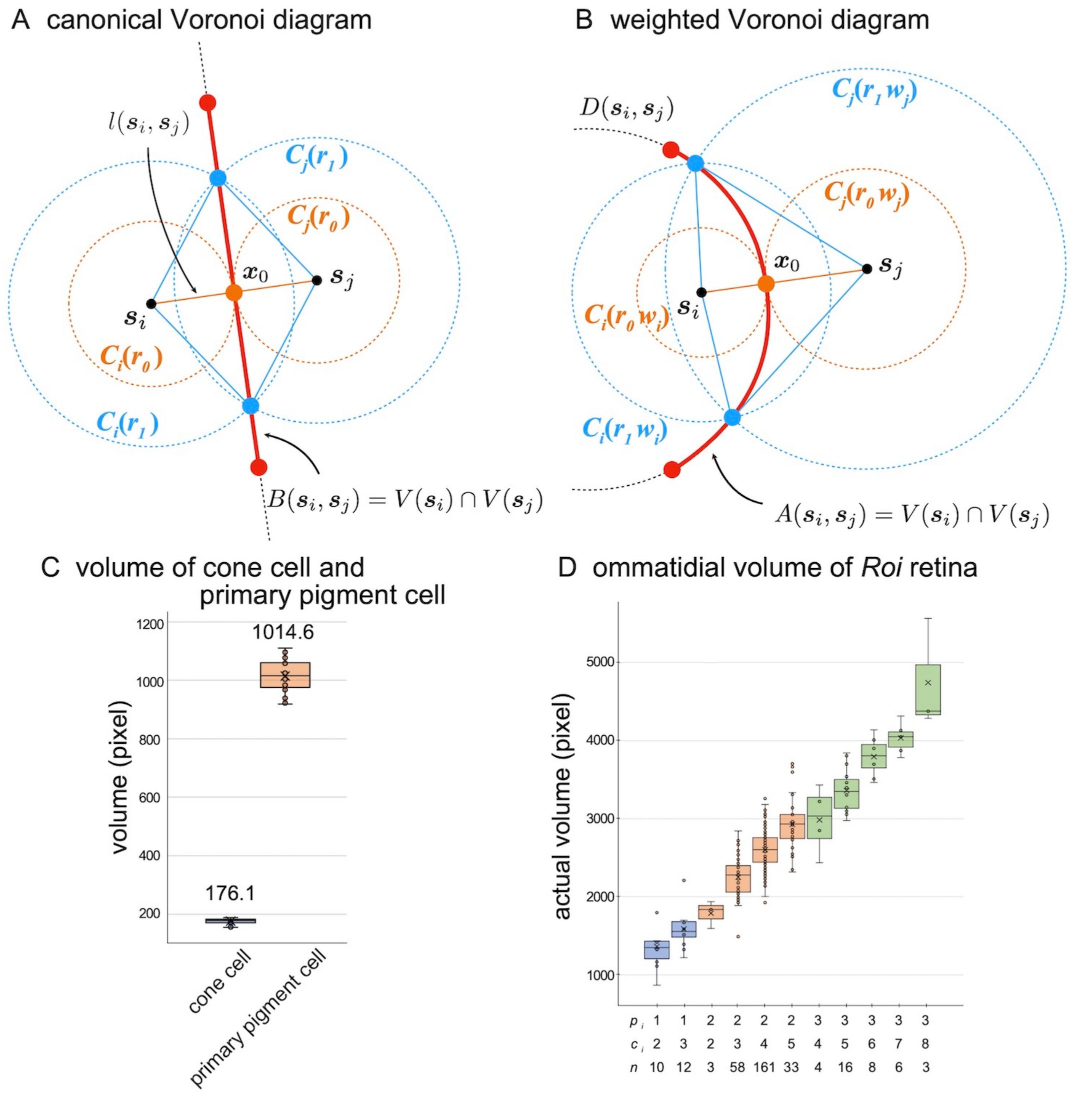


**Fig. S5. Definition of canonical and weighted Voronoi diagrams.**

(A) In the canonical Voronoi diagram, a part of the vertical bisectors, $l(\boldsymbol{s}_i, \boldsymbol{s}_j)$, between two mother points $\boldsymbol{s}_i$ and $\boldsymbol{s}_j$, constitutes the Voronoi edge, $B(\boldsymbol{s}_i, \boldsymbol{s}_j)$. $C_i(r)$ and $C_j(r)$ are the circles of radius $r$ emanating from $\boldsymbol{s}_i$ and $\boldsymbol{s}_j$. respectively, and intersect on $B(\boldsymbol{s}_i, \boldsymbol{s}_j)$. (B) In the weighted Voronoi diagram, $w_i$ and $w_j$ are the relative growth rates of the circles $C_i(rw_i)$ and $C_j(rw_j)$ emanating from $\boldsymbol{s}_i$ and $\boldsymbol{s}_j$. respectively. $C_i(rw_i)$ and $C_j(rw_j)$ intersect on the Voronoi edge $A(\boldsymbol{s}_i, \boldsymbol{s}_j)$, which is a part of the circle $D(\boldsymbol{s}_i, \boldsymbol{s}_j)$. (C) Volumes of individual cone and primary pigment cells in control ommatidia (n=36). (D) Ommatidia from five compound eyes are classified according to the number of primary pigment cells ($p_i$) and cone cells ($c_i$), and their actual volumes are plotted. $n$ indicates the number of ommatidia for each class. Cross, mean; center line, median; box limits, upper and lower quartiles; whiskers, 1.5x interquartile range (C, D).
